# Dynamics of the ACE2 - SARS-CoV/SARS-CoV-2 spike protein interface reveal unique mechanisms

**DOI:** 10.1101/2020.06.10.143990

**Authors:** Amanat Ali, Ranjit Vijayan

## Abstract

The coronavirus disease 2019 (COVID-19) pandemic, caused by the severe acute respiratory syndrome coronavirus 2 (SARS-CoV-2), is a major public health concern. A handful of static structures now provide molecular insights into how SARS-CoV-2 and SARS-CoV interact with its host target, which is the angiotensin converting enzyme 2 (ACE2). Molecular recognition, binding and function are dynamic processes. To evaluate this, multiple all atom molecular dynamics simulations of at least 500 ns each were performed to better understand the structural stability and interfacial interactions between the receptor binding domain of the spike protein of SARS-CoV-2 and SARS-CoV bound to ACE2. Several contacts were observed to form, break and reform in the interface during the simulations. Our results indicate that SARS-CoV and SARS-CoV-2 utilizes unique strategies to achieve stable binding to ACE2. Several differences were observed between the residues of SARS-CoV-2 and SARS-CoV that consistently interacted with ACE2. Notably, a stable salt bridge between Lys417 of SARS-CoV-2 spike protein and Asp30 of ACE2 as well as three stable hydrogen bonds between Tyr449, Gln493, and Gln498 of SARS-CoV-2 and Asp38, Glu35, and Lys353 of ACE2 were observed, which were absent in the SARS-CoV-ACE2 interface. Some previously reported residues, which were suggested to enhance the binding affinity of SARS-CoV-2, were not observed to form stable interactions in these simulations. Stable binding to the host receptor is crucial for virus entry. Therefore, special consideration should be given to these stable interactions while designing potential drugs and treatment modalities to target or disrupt this interface.

## Introduction

The recent outbreak of coronavirus disease 2019 (COVID-19), caused by the novel severe acute respiratory syndrome coronavirus 2 (SARS-CoV-2), has affected all walks of life. Genomic studies have established that SARS-CoV-2 belong to the betacoronavirus genus, which also includes SARS-CoV and MERS-CoV that were associated with previous outbreaks of relatively smaller scale ^1–3^. These coronaviruses attach to the host cell with the aid of the spike glycoprotein present on its envelope. Coronavirus spike glycoprotein is composed of two subunits - the S1 subunit is important for binding to the host cell receptor and the S2 subunit is responsible for the fusion of the virus and the host cell’s membrane. Angiotensin converting enzyme 2 (ACE2), an enzyme located on the outer surface of a wide variety of cells, is the primary host cell target with which the spike protein of SARS-CoV and SARS-CoV-2 associates ^4–6^. The receptor binding domain (RBD) of the S1 subunit of these viruses binds to outer surface of the claw like structure of ACE2^7^.

The sequence similarity of the RBD region of SARS-CoV and SARS-CoV-2 spike protein is between 73% to 76% ^7^. Fourteen residues of the SARS-CoV spike protein RBD have been reported to interact with human ACE2. These are Tyr436, Tyr440, Tyr442, Leu443, Leu472, Asn473, Tyr475, Asn479, Gly482, Tyr484, Thr486, Thr487, Gly488, and Tyr491 ^8,9^. Only eight of these residues are conserved in SARS-CoV-2 ^9^ The equivalent conserved residues in SARS-CoV-2 are Tyr449, Tyr453, Asn487, Tyr489, Gly496, Thr500, Gly502, and Tyr491, while Leu455, Phe456, Phe486, Gln493, Gln498, and Asn501 are substituted.

The atomic level three-dimensional structure of several SARS-CoV-2 proteins have now been determined ^10–13^. These X-ray crystallography based structures provide remarkable insights into macromolecular structure and intermolecular interactions. However, molecular recognition and binding are dynamic processes. Molecular dynamics (MD) simulation often complement traditional structural studies for looking at the dynamics of these processes at the atomic level ^14,15^. Such simulations can provide insights into the structural stability of macromolecular complexes, flexibility of interacting subunits and the interactions of residues in the interface.

Here, we report the stability and binding dynamics of SARS-CoV-2 spike RBD bound to ACE2, and compare this with the dynamics of SARS-CoV spike RBD, by running multiple 500 ns all-atom MD simulations. High resolution X-ray crystal structures of SARS-CoV-2 spike RBD bound to ACE2 illustrate fourteen positions that are associated with the interaction between SARS-CoV/SARS-CoV-2 and ACE2. The primary objective of this study was to identify both similarities and dissimilarities in the dynamics of the interactions between SARS-CoV/SARS-CoV-2 and ACE2 and to identify key residues that could be vital to the integrity of this interface. This would provide insights into residues that could be targeted for disrupting this interface.

## Results

### MD simulations of SARS-CoV-2-ACE2 and SARS-CoV-ACE2 complexes

500 ns MD simulations of SARS-CoV-ACE2 complex (PDB ID: 6M0J) and SARS-CoV-ACE2 complex (PDB: 2AJF) were performed in triplicate. One of the 500 ns simulations of SARS-CoV-2-ACE2 complex was further extended to 1 μs to ensure that the interactions are faithfully retained for a longer duration. Additionally, triplicate 500 ns simulations of a second SARS-CoV-2-ACE2 structure (PDB ID: 6LZG) was also performed to ensure that results were not biased by a single structure. The overall structural integrity of all stimulations of SARS-CoV2-ACE2 complex were retained with a Cα root mean square deviation (RMSD) from the starting structure that was less than 4 Å. In the case of the SARS-CoV-ACE2 simulations, one of simulations had a Cα RMSD that was under at 5 Å. The oscillating RMSD of this simulation was characteristic of the closing/opening motion of the ACE2 claw like structure ^16^ (Figures 1A and Figure 1B). The other two SARS-CoV-ACE2 simulations had a higher RMSD that was associated with the detachment of the spike protein from one end of interface (Figure 1B). Interestingly, the other end remained attached as evident from specific contacts that were retained throughout the simulations. Protein secondary structure composition and compactness, as indicated by the radius of gyration, of ACE2 and spike protein structures were also preserved throughout the simulations (Supplementary Figure 1).

**Figure 1.**
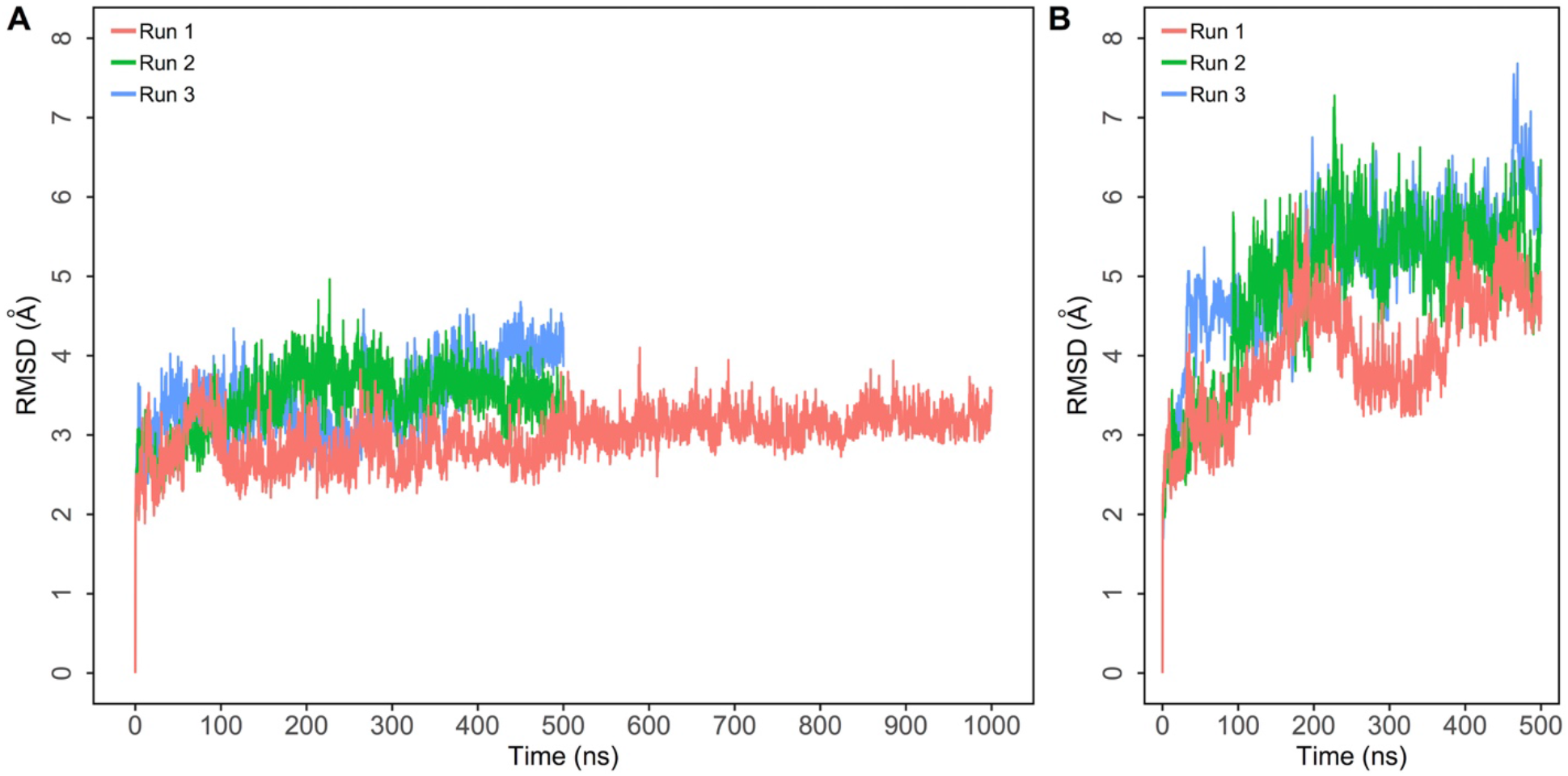
Root mean square deviation (RMSD) of protein Cα atoms with respect to the initial structure obtained from three independent 500 ns runs of SARS-CoV-2-ACE2 and SARS-CoV bound simulations of ACE2. The first SARS-CoV-2-ACE2 simulation was extended to 1 μs. A) Simulations of SARS-CoV-2-ACE2 complex; B) Simulations of SARS-CoV-ACE2 complex.

### Comparison of regional fluctuations in the SARS-CoV-2-ACE2 and SARS-CoV-ACE2 complexes

To identify and compare backbone stability and fluctuations of the two complexes, root mean square fluctuation (RMSF) of backbone Cα atoms were computed and plotted (Figure 2). This was also projected as beta factors in PDB structures and visualized (Supplementary Figure 2). Three loops (residues 474-485, 488-490, and 494-505) of the SARS-CoV-2 spike protein RBD make contact with ACE2. The homologous region in SARS-CoV range between 461-471, 474-476, and 480-491. In the SARS-CoV2-ACE2 complex, the loops between residues 474-485 and 488-490 exhibited a comparatively higher fluctuation with respect to the rest of the RBD structure while the 494-505 loop demonstrated very limited fluctuations (Figure 2B) However, in SARS-CoV RBD, both loops demonstrated higher fluctuations throughout the simulations (Figure 2D). Interestingly, the region around Lys417 showed very limited fluctuations in SARS-CoV-2. Overall, the backbone of SARS-CoV-2 exhibited lower fluctuations compared to SARS-CoV (Figures 2B and Figure 2D).

**Figure 2.**
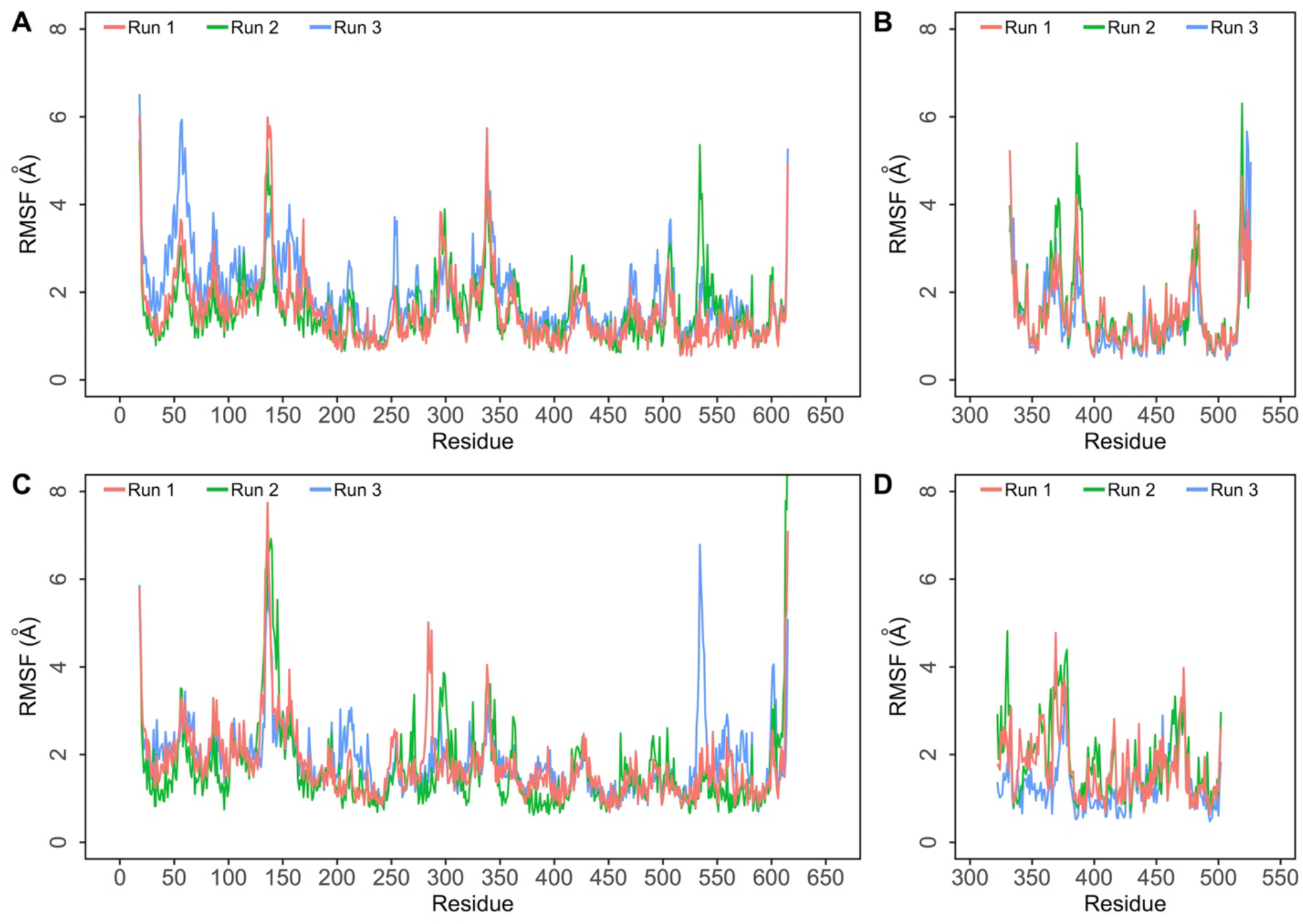
mean square fluctuation (RMSF) of protein Cα atoms obtained from three independent runs of SARS-CoV-2 and SARS-CoV bound simulations of ACE2. A) RMSF of Cα atoms of ACE2 protein in the SARS-CoV-2-ACE2 complex; B) RMSF of Cα atoms of SARS-CoV-2 spike RBD in the SARS-CoV-2-ACE2 complex; C) RMSF of Cα atoms of ACE2 protein in the SARS-CoV-ACE2 complex; D) RMSF of Cα atoms of SARS-CoV spike RBD in the SARS-CoV-2-ACE2 complex.

Spike RBD structures were extracted from the SARS-CoV-2 and SARS-CoV complexes and simulated independently to study how the RBD structures behave when not bound to ACE2. The RMSF of the RBD-only structures were then compared to the corresponding RBD bound to ACE2. Interestingly, in the SARS-CoV-2 RBD, residues between 469-505, that form loops, exhibited higher fluctuations when not bound to ACE2 (Supplementary Figure 3A). The equivalent region of the SARS-CoV RBD did not produce such a marked difference in bound and unbound simulations (Supplementary Figure 3B). It would appear that this region in SARS-CoV-2 RBD is effectively stabilized after binding to ACE2. Several residues between 484-505 are vital to the ACE2-spike RBD interface. The loop region between residues 384-392 in SARS-CoV-2 fluctuated more when bound to ACE2, while there was no noticeable difference in the SARS-CoV simulations (Supplementary Figures 3A and 3B). This loop is not located in the interface and, thus, it is not clear if it has any significance.

To look at the effect of SARS-CoV2 and SARS-CoV binding on backbone stability and residue fluctuations of ACE2, the root mean square fluctuation (RMSF) of backbone Cα atoms of ACE2 were evaluated (Figures 2A and 2C). The fluctuation of ACE2 backbone in both SARS-CoV-2 and SARS-CoV complexes were comparable and showed limited fluctuations except in loop regions (residues 130-138 and 278-291) where the SARS-CoV-2 bound complex showed lower fluctuations compared to SARS-CoV bound complex (Figures 2A and 2C). More importantly, residues in the interfacial region of ACE2 (residues 78-83 and 353-357) exhibited slightly lower fluctuations in SARS-CoV-2. Additionally, loop regions of ACE2 - 82-89 and 351-354 - also exhibited slightly lower fluctuations in SARS-CoV-2 (Figures 2A and 2C).

### Interfacial residue contact duration differs substantially between SARS-CoV-2 and SARS-CoV bound complexes

Several intermolecular contacts were observed to form, break and reform during the simulations. This included salt bridges, hydrogen bonds, *π-π* and cation-*π* interactions (Supplementary Table 1). Some of these interactions were more persistent than others. The residues of SARS-CoV-2/SARS-CoV spike RBD that interacted with ACE2 consistently are shown in Figures 3A and 3B. The duration of specific intermolecular contacts between SARS-CoV-2/SARS-CoV and ACE2 interfaces and the dynamics of the salient ones along the length of the simulation trajectories are shown in Figure 4 and Supplementary Figures 4 and 5.

**Figure 3.**
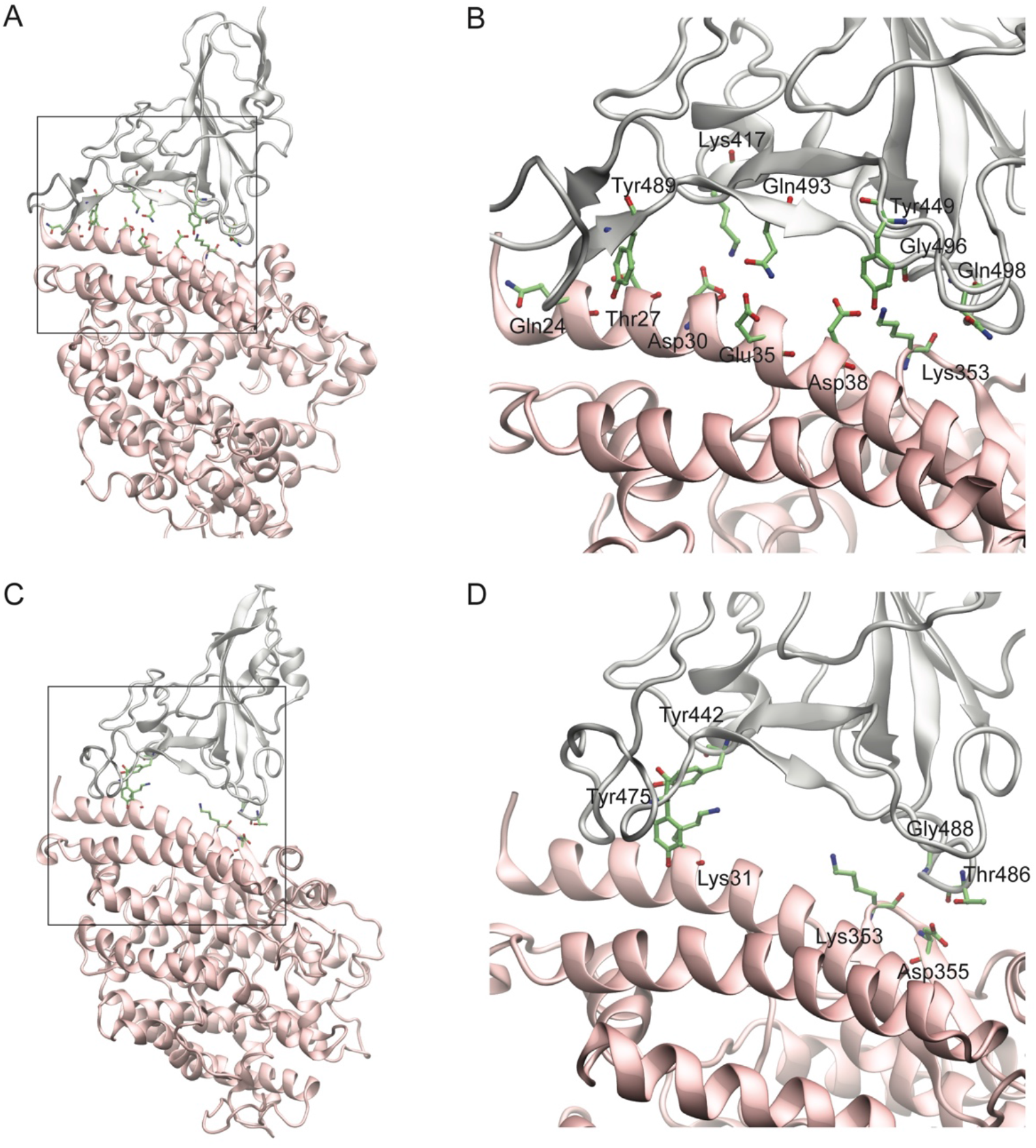
Structures of the SARS-CoV-2 and SARS-CoV spike RBD (gray) bound to ACE2 (pink). The interface is enlarged in the adjacent image showing the interacting residues in green stick representation. A) SARS-CoV-2 spike protein RBD (gray) bound to ACE2 (pink); B) The boxed region in A is enlarged showing the residues that interact in the interface. C) SARS-CoV spike protein RBD (gray) bound to ACE2 (pink); D) The boxed region in C is enlarged showing the residues that interact in the interface.

**Figure 4.**
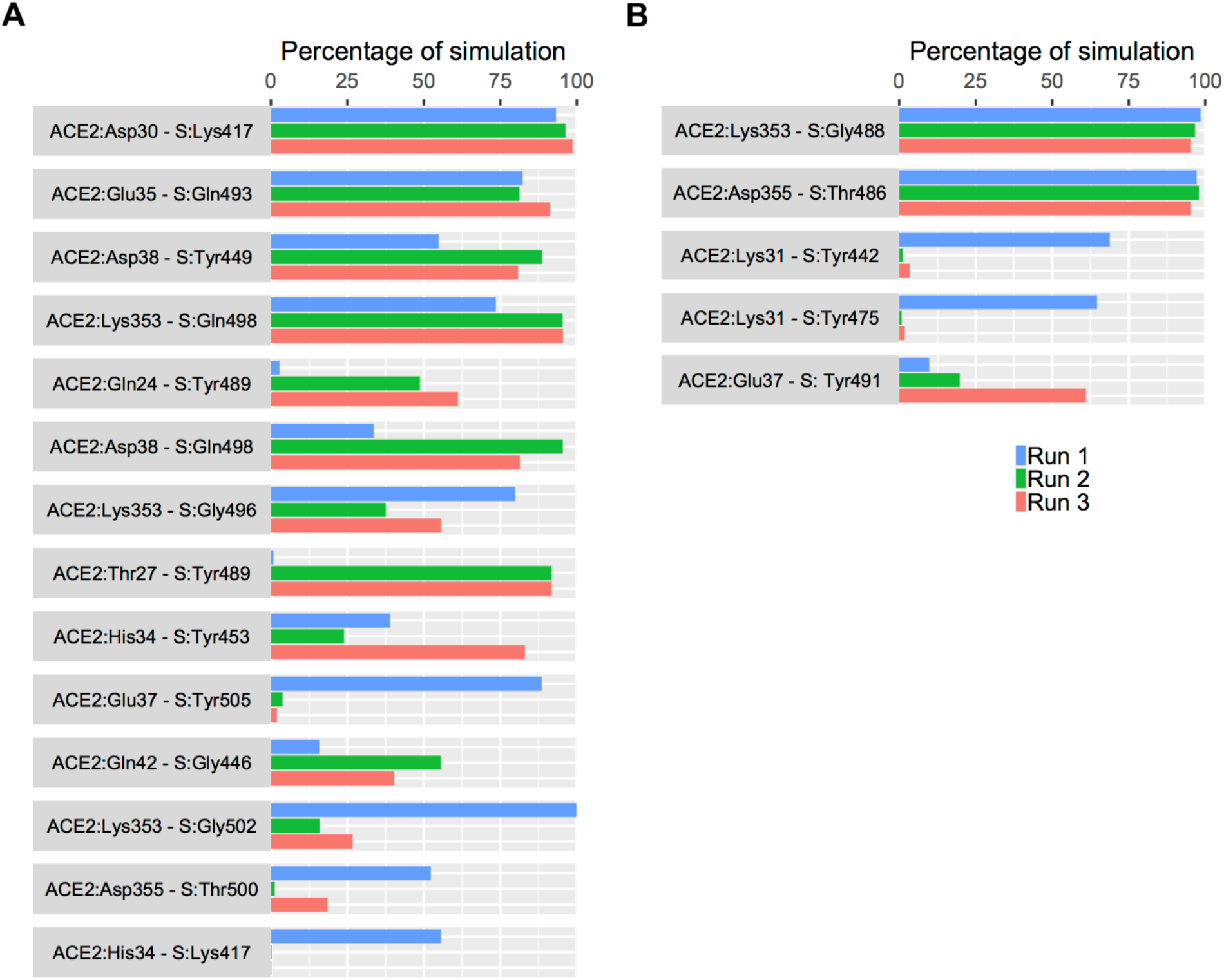
The percentage of simulation time during which intermolecular contacts were retained between ACE2 and SARS-CoV-2/SARS-CoV spike protein RBD residues. A) Data from three independent runs of the SARS-CoV-2-ACE2 complex B) Data from three independent runs of the SARS-CoV-ACE2 complex.

Lys31 and Lys353 of ACE2 normally forms intramolecular salt bridges with Glu35 and Asp38 and are buried in a hydrophobic environment ^4,8^. In the SARS-CoV simulation where the spike RBD remained stably bound to ACE2, the Lys31 hot spot was observed to consistently form cation-π interactions with Tyr442 and Tyr475 of SARS-CoV spike RBD in only one simulation (Figure 4B). However, Leu455 and Tyr489, the residues at equivalent positions in SARS-CoV-2 did not interact with Lys31 in the SARS-CoV-2 simulations. Instead, Glu484 and Gln493 of SARS-CoV-2 formed extremely weak interactions with Lys31 (Supplementary Table 1). The likelihood of an interaction between Gln493 of SARS-CoV-2 and Lys31 residue of ACE2 was reported recently ^7^.

Next, the backbone of Gly488 of SARS-CoV consistently interacted with Lys353 of ACE2 in all three simulations, while the Gly502 at the equivalent location in SARS-CoV-2 formed sustained interactions with Lys353 of ACE2 in only one simulation (Figures 4A and 4B). Additionally, the side chain of Gln498 and backbone of Gly496 of SARS-CoV-2 spike RBD formed sustained interactions in three and two simulations, respectively, with Lys353 of ACE2. Such an interaction was absent in equivalent residues of SARS-CoV (Figures 4A and 4B). This suggests that SARS-CoV-2 and SARS-CoV differs in how they target the basic Lys31 and Lys353 residues of ACE2.

Tyr449 and Tyr489, two conserved residues in SARS-CoV-2 spike RBD, consistently interacted with Asp38 and Gln24 of ACE2 in multiple simulations. Tyr453 and Tyr505 were observed to interact with His34 and Gln37, respectively, of ACE2 for at least 80% of one simulation each (Figures 4A and 4B). The corresponding residues in SARS-CoV did not appear to form such sustained interactions except Tyr491 which consistently interacted with Gln37 in two simulations. Furthermore, Gln493 and Gln498 of SARS-CoV-2 showed sustained interactions with Glu35 and Asp38, respectively, while the corresponding Asn479 and Tyr484 in SARS-CoV exhibited extremely weak interactions (Figures 4A and 4B). This is in agreement with recent work that shows the likely existence of Tyr449-Asp38, Tyr453-His34, Tyr489-Gln24, and Gln493-Glu35 interactions between SARS-CoV-2 spike RBD and ACE2 ^16–18^.

Three residues which are mutated in SARS-CoV-2 (Ala475, Lys417 and Gly446) interacted with ACE2 residues (Gln24, Asp30 and Gln42). Such interactions were not observed in corresponding residues of SARS-CoV (Figures 4A and 4B, Supplementary Table 1). Significantly, a very strong salt bridge was established and sustained between Lys417 of SARS-CoV-2 spike RBD and Asp30 of ACE2 for nearly the full duration of all simulations. Notably, this salt bridge is absent in SARS-CoV since the equivalent residue is Val404, which is incapable of forming such an interaction (Figures 4A and 4B). Gly446 maintained an interaction with Gln42 of ACE2 for majority of only one simulation (Figure 4A), while Ala475 exhibited only weak interactions with Gln24 in the simulations.

To look at the dynamics of the interface, interactions that were maintained for at least 50% of the total simulation time in three simulations in the two complexes were evaluated. Four interfacial residues in SARS-CoV-2 (Lys417, Gln493, Tyr449 and Gln498) and two in SARS-CoV (Thr486 and Gly488) maintained such interactions with four (Asp30, Glu35, Asp38 and Lys353) and two (Lys353 and Asp355) residues of ACE2 respectively (Figures 4A and 4B). Hence, there are noteworthy differences between how the two viral spike proteins interact with ACE2 and the larger number of sustained interactions in SARS-CoV-2 spike protein could be associated with a higher binding affinity of SARS-CoV-2. Additionally, similar dynamics of interacting residues were also observed in triplicate 500 ns simulations of another complex structure of SARS-CoV-2-ACE2 (PDB: 6LZG) that was simulated to ensure that the results were not biased by one structure (Supplementary Figure 6 and Supplementary Table 1).

The total number of intermolecular hydrogen bonds between the two complexes were monitored throughout the simulation. A higher number of hydrogen bonds were observed in the SARS-CoV-2-ACE2 complex when compared to SARS-CoV-ACE2 complex (Supplementary Figure 7). Contact distance of the most stable interfacial residues of SARS-CoV-2 and SARS-CoV were also evaluated and plotted since this gives a better indication of interactions rather than preset cutoffs used for determination of an interaction. The contact distance density plot of consistently interacting residues showed sharper peaks (Supplementary Figures 4 and 5). The fewer interactions and hydrogen bonds in the second and third simulation of the SARS-CoV-2 bound complex was due to the notable separation of the RBD from one end of the interface near the Lys31 residue of ACE2 (Figures 3 and 4 and Supplementary Figure 2).

Recent studies, based on models of the SARS-CoV-2 spike protein RBD, have indicated that Leu455, Phe486, Ser494, and Asn501 of SARS-CoV-2 are important for binding to ACE2 via their interaction with Met82, Tyr83, Lys31, and Tyr41 residues ^3,7,18^. However, from the simulations, only a weak π-π interaction was observed between Phe486 and Tyr83 (Supplementary Table 1). While, Leu455, Ser494, and Asn501 were not observed to form any significant interactions with ACE2.

Water molecules play an important role in many intermolecular interfaces. In this instance, six conserved water sites were found in the interface between ACE2 and SARS-CoV-2 spike RBD. Water-mediated indirect interactions were formed between ACE2:Lys31 and spike:Phe490/Leu492, ACE2:Asp38 and spike:Gly496, ACE2:Asn33/His34/Glu37/Asp38 and spike:Arg403 (Figure 5). These could also play a role in stabilizing the interface.

**Figure 5.**
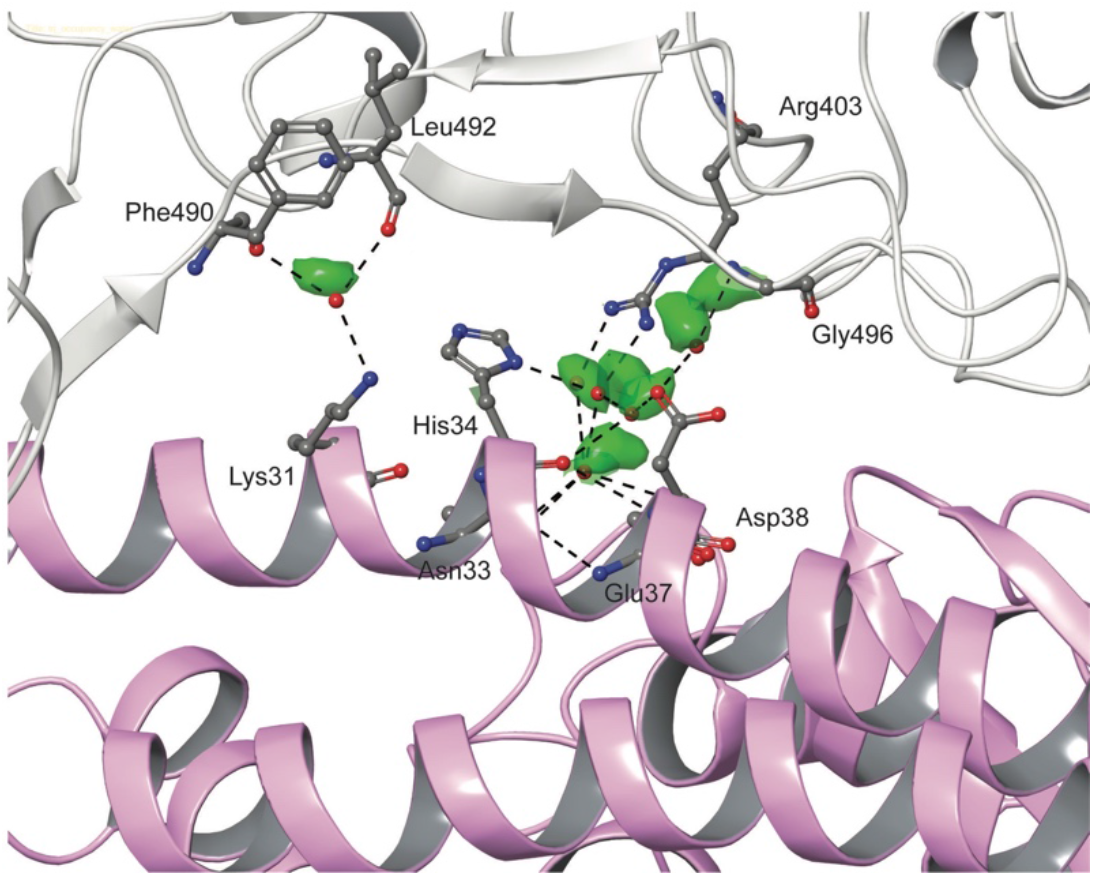
Conserved water sites in the interface between SARS-CoV-2 spike RBD (gray) and ACE2 (pink) obtained from a 1 μs MD simulation of the complex. Regions which high residence time (green) were identified by constructing a 3D histogram of water occupancy sites. Specific water molecules from a frame of the simulation are shown as red spheres. For clarity only the oxygen atom of water molecules is shown. The interacting residues are shown in stick representation and hydrogen bonds are shown with black dashed lines.

## Discussion

This study provides insight into the stability of the interactions that define the SARS-CoV-ACE2 and SARS-CoV-2-ACE2 interfaces, using extended MD simulations of X-ray crystal structures of these complexes. Firstly, interactions that were shared by SARS-CoV-2-ACE2 and SARS-CoV-ACE2 complexes were assessed. SARS-CoV-2 spike protein RBD consistently interacted with ACE2 in three clusters. At one end, Gln493 and Gln498 formed sustained hydrogen bonds with Glu35 and Lys353 of ACE2. At the other end, Tyr449 formed hydrogen bond with Asp38 of ACE2. In the middle, Lys417 formed a strong and stable salt bridge with Asp30 of ACE2. Additionally, several intermittent interactions of SARS-CoV-2 could permit it to bind more stably than SARS-CoV (Figures 3B and 4A and Supplementary Table 1). These findings were also supported by the results from two different structures of the SARS-CoV-2-ACE2 complex (Supplementary Figure 6 and Supplementary Table 1). Loop regions (residues 484-505) of SARS-CoV-2 fluctuated less when bound to ACE2, when compared to SARS-CoV. This is stabilized by the formation of sustained interactions between Gln493 and Gln498 of SARS-CoV-2 and Glu35 and Lys353 of ACE2, respectively (Figures 4A and Supplementary Figure 4).

Unlike SARS-CoV-2, in SARS-CoV, the region in the middle was devoid of any stable interactions with ACE2. However, at the two ends, a different set of residues in SARS-CoV formed interactions with ACE2 (Figure 3D). Therefore, it is apparent that there are several similarities and differences in the structure and dynamics of the interactions of SARS-CoV-2 and SARS-CoV with ACE2. Hence, antibodies or antiviral treatment modalities that target the spike protein of SARS-CoV is not expected to produce a similar effect with SARS-CoV-2. Some of the recent studies that failed to inhibit the binding of SARS-CoV-2 RBD to ACE2, support these findings ^13,19^.

Secondly, two charged virus binding hot spots on human ACE2 (Lys31 and Lys353), which are essential for the binding of SARS-CoV, have been studied extensively ^4,8^. Charge neutralization of these hot spot lysines has been shown to be important for the binding of coronavirus to ACE2 ^7,8^. SARS-CoV-2 and SARS-CoV utilizes unique strategies to achieve this. Interestingly, SARS-CoV-2 residues only formed sustained interactions with Lys353 of ACE2 and these were absent in SARS-CoV indicating an adaptation to a stronger interface (Figure 4A and 4B). Several of these vital residues fall in loop regions of SARS-CoV-2 and SARS-CoV. Three loop regions - 474-485, 488-490, and 494-505 of SARS-CoV-2 demonstrated limited fluctuation compared to the corresponding region in SARS-CoV (Figure 2B and 2D). Lower fluctuations, observed in residues of SARS-CoV-2 that bind to the Lys353 hot spot of ACE2, could be another reason for the better affinity of SARS-CoV-2 to ACE2 ^7,18^.

Thirdly, the non-conserved residue Lys417 of SARS-CoV-2 formed a very stable salt bridge with Asp30 of ACE2. This interaction was absent in the corresponding residue (Val404) of SARS-CoV (Figure 4A and 4B). Lys417 provides a positively charged patch on the RBD of SARS-CoV-2 which is absent in SARS-CoV, where the corresponding residue is Val404. The RBD is also stabilized by this strong interaction; the RMSF of the region around Lys417 demonstrated lower fluctuations in SARS-CoV-2 RBD compared to the corresponding region of SARS-CoV (Figure 2B and 2D). The strength of this sustained salt bridge between Lys417 and Asp30 could contribute to the substantially different binding affinity of SARS-CoV-2 to ACE2 receptor when compared to SARS-CoV ^16,18^. Notably, some of the previously reported residues (Leu455, Phe456, Tyr473, Phe486, Ser494, and Asn501) that were suggested to enhance the binding affinity of SARS-CoV-2, were not observed in these simulations ^3,16,18^.

In conclusion, while SARS-CoV-2 and SARS-CoV spike RBD bind to the same region of ACE2 and share several similarities in how they interact with ACE2, there are a number of differences in the dynamics of the interactions. One salient difference is the presence of a stable salt bridge between Lys417 of SARS-CoV-2 spike protein and Asp30 of ACE2 as well as three stable hydrogen bonds between Tyr449, Gln493, and Gln498 of SARS-CoV-2 and Asp38, Glu35, and Lys353 of ACE2, which were not observed in the SARS-CoV-ACE2 interface. Stable viral binding with the host receptor is crucial for virus entry. Thus, special consideration should be given to these stable interactions while designing potential drugs and treatment modalities to target or disrupt this interface.

## Materials and methods

Coordinates of the three dimensional X-ray crystal structures of the SARS-CoV and SARS-CoV-2 RBD in complex with ACE2 were obtained from the Protein Data Bank (PDB). The PDB IDs of the structure used are 6M0J and 6LZG for SARS-CoV-2 spike protein RBD bound to ACE2 and 2AJF for the SARS-CoV spike protein RBD bound to ACE2. Schrödinger Maestro 2019-4 (Schrödinger, LLC, New York, NY) was used to visualize and prepare the protein structures for simulations. The structures were first pre-processed using the Protein Preparation Wizard (Schrödinger, LLC, New York, NY). The protein preparation stage included proper assignment of bond order, adjustment of ionization states, orientation of disorientated groups, creation of disulfide bonds, removal of unwanted water molecules, metal and co-factors, capping of the termini, assignment of partial charges, and addition of missing atoms and side chains. In the case of the SARS-CoV structure, a loop (residues 376-381) missing in the PDB structure was modeled using Schrödinger Prime ^20^. Hydrogen atoms were incorporated, and standard protonation state at pH 7 was used. Structures of spike protein RBD bound to ACE2 were placed in orthorhombic boxes of size 125 Å × 125 Å × 125 Å and solvated with single point charge (SPC) water molecules using the Desmond System Builder (Schrödinger, LLC, New York, NY). A box size of 85 Å × 85 Å × 85 Å was used for simulations of spike RBD structures isolated from the PDB structures 6M0J and 2AJF. Simulation systems were neutralized with counterions and a salt concentration of 0.15M NaCl was maintained. MD simulations were performed using Desmond ^21^. The OPLS forcefield was used for all calculations. All systems were subjected to Desmond’s default eight stage relaxation protocol before the start of the production run. 500 ns simulations were performed in triplicate with a different set of initial velocities for simulations involving 6M0J, 6LZG and 2AJF while one 500 ns simulation each were run for the isolated spike protein structures. One of the 500 ns simulations of the SARS-CoV-2-ACE2 complex was extended to 1 μs to ensure that the interactions are retained for a longer period. For the simulations, the isotropic Martyna-Tobias-Klein barostat and the Nose-Hoover thermostat were used to maintain the pressure at 1 atm and temperature at 300 K, respectively ^22,23^. Short-range cutoff was set as 9.0 Å and long-range coulombic interactions were evaluated using the smooth particle mesh Ewald method (PME) ^24^. A time-reversible reference system propagator algorithm (RESPA) integrator was employed with an inner time step of 2.0 fs and an outer time step 6.0 fs ^25^. Simulation data was analyzed using packaged and in house scripts and plotted using R 3.6.0 (https://www.r-project.org).

## Acknowledgment

This study was supported by a UPAR grant (31S243) from United Arab Emirates University to RV. The funder had no role in the study design, data collection and interpretation, or the decision to submit the work for publication.

## Author contributions

RV conceived the idea and performed the experiments. RV, AA analyzed and wrote the manuscript.

