## Supplementary Information for "Dynamics of the ACE2 - SARS-CoV/SARS-CoV-2 spike protein interface reveal unique mechanisms"

**Table 1**. Percentage contact time of equivalent interacting residues of SARS-CoV-2 and SARS-CoV spike RBD with ACE2.

| ACE2 residue | SARS-CoV-2 spike protein residue | Percentage contact time (PDB ID: 6M0J) | | | Percentage contact time (PDB ID: 6LZG) | | | SARS-CoV spike protein residue | Percentage contact time (PDB ID: 2AJF) | | |
|---|---|---|---|---|---|---|---|---|---|---|---|
| | | Run 1 | Run 2 | Run 3 | Run 1 | Run 2 | Run 3 | | Run 1 | Run 2 | Run 3 |
| **Asp30** | **Lys417** | 93.2 | 96.3 | 98.6 | 98.0 | 92.8 | 86.4 | **Val404** | Nil | Nil | Nil |
| **Glu35** | **Gln493** | 82.3 | 81.3 | 91.2 | 90.3 | 86.0 | 84.6 | **Asn479** | 11.7 | 11.5 | 16.4 |
| **Asp38** | **Tyr449** | 54.8 | 88.6 | 80.8 | 10.0 | 13.2 | 45.9 | **Tyr436** | 10.0 | 6.5 | 2.7 |
| **Lys353** | **Gln498** | 73.5 | 95.3 | 95.5 | 90.7 | 22.3 | 90.7 | **Tyr484** | 1.0 | 0.6 | 0.5 |
| **Gln24** | **Tyr489** | 2.8 | 48.7 | 61.1 | 0.04 | 24.5 | 22.2 | **Tyr475** | Nil | Nil | Nil |
| **Asp38** | **Gln498** | 33.6 | 95.4 | 81.5 | 84.6 | 57.4 | 95.6 | **Tyr484** | 2.0 | 4.1 | 0.4 |
| **Lys353** | **Gly496** | 79.9 | 37.6 | 55.6 | 83.6 | 11.6 | 28.8 | **Gly482** | 12.0 | 4.4 | 1.8 |
| **Thr27** | **Tyr489** | 0.77 | 91.8 | 91.8 | 0.06 | 0.02 | 13.2 | **Tyr475** | Nil | 0.2 | Nil |
| **His34** | **Tyr453** | 39.0 | 23.9 | 83.1 | 36.1 | 29.4 | 12.6 | **Tyr440** | 0.01 | Nil | 20.3 |
| **Glu37** | **Tyr505** | 88.5 | 3.8 | 1.1 | 71.0 | 36.1 | 6.6 | **Tyr491** | 20.0 | 64.7 | 69.3 |
| **Gln42** | **Gly446** | 15.8 | 55.5 | 40.3 | 0.04 | 10.1 | 37.7 | **Thr433** | 0.15 | 2.5 | Nil |
| **Lys353** | **Gly502** | 99.9 | 16.0 | 26.7 | 99.8 | 6.7 | 2.4 | **Gly488** | 99.0 | 99.0 | 97.0 |
| **Asp355** | **Thr500** | 52.3 | 1.6 | 18.5 | 56.9 | 4.7 | 1.5 | **Thr486** | 97.0 | 99.4 | 96.1 |
| **His34** | **Lys417** | 55.6 | 0.23 | 0.2 | 34.9 | 54.6 | 25.2 | **Val404** | Nil | Nil | Nil |
| **Lys31** | **Leu455** | Nil | Nil | Nil | Nil | Nil | Nil | **Tyr442** | 68.8 | 7.8 | 2.0 |
| **Tyr83** | **Phe486** | 8.6 | 26.8 | 29.2 | 17.9 | 25.6 | 29.1 | **Leu472** | Nil | Nil | Nil |
| **Lys31** | **Tyr489** | 0.3 | 0.02 | Nil | 2.1 | 0.2 | 0.2 | **Tyr475** | 64.7 | 13.8 | 1.9 |
| **Gln24** | **Ala475** | 30.6 | 18.4 | 35.7 | 27.5 | 10.7 | 10.6 | **Pro462** | Nil | 0.2 | Nil |

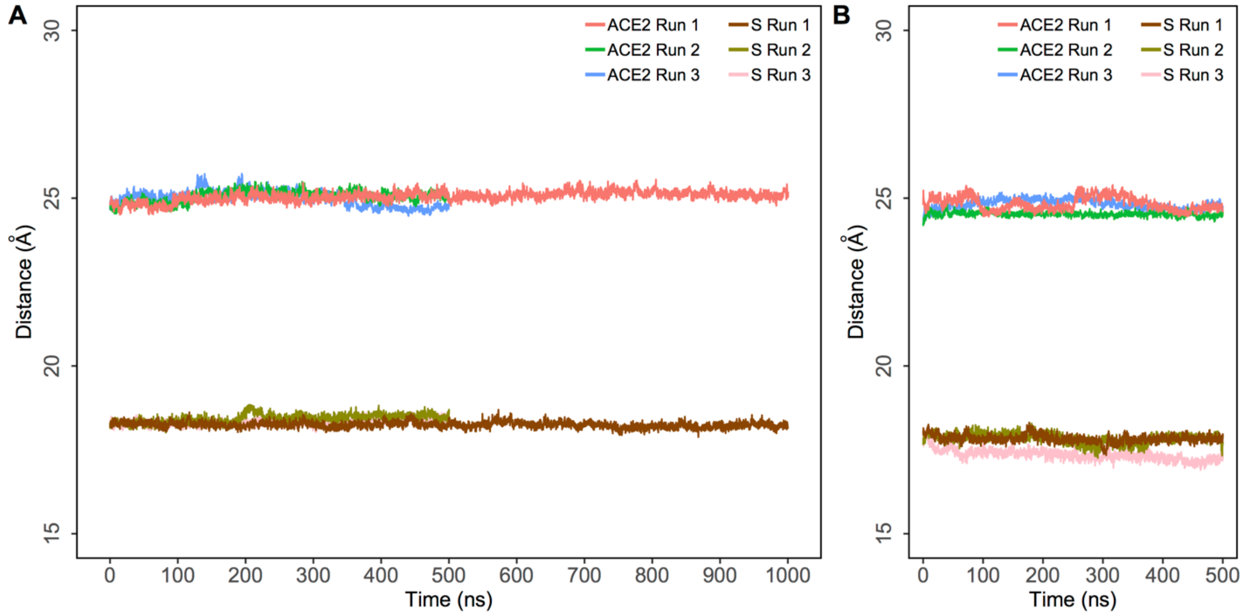


**Figure 1.** Radius of gyration (Rg) of ACE2 and spike protein structures obtained from three independent simulations. All simulations were run for 500 ns while the first simulation of the ACE2-SARS-CoV-2 complex was extended to 1 μs. A) Rg of ACE2-SARS-CoV-2 complex; B) Rg of ACE2-SARS-CoV complex.

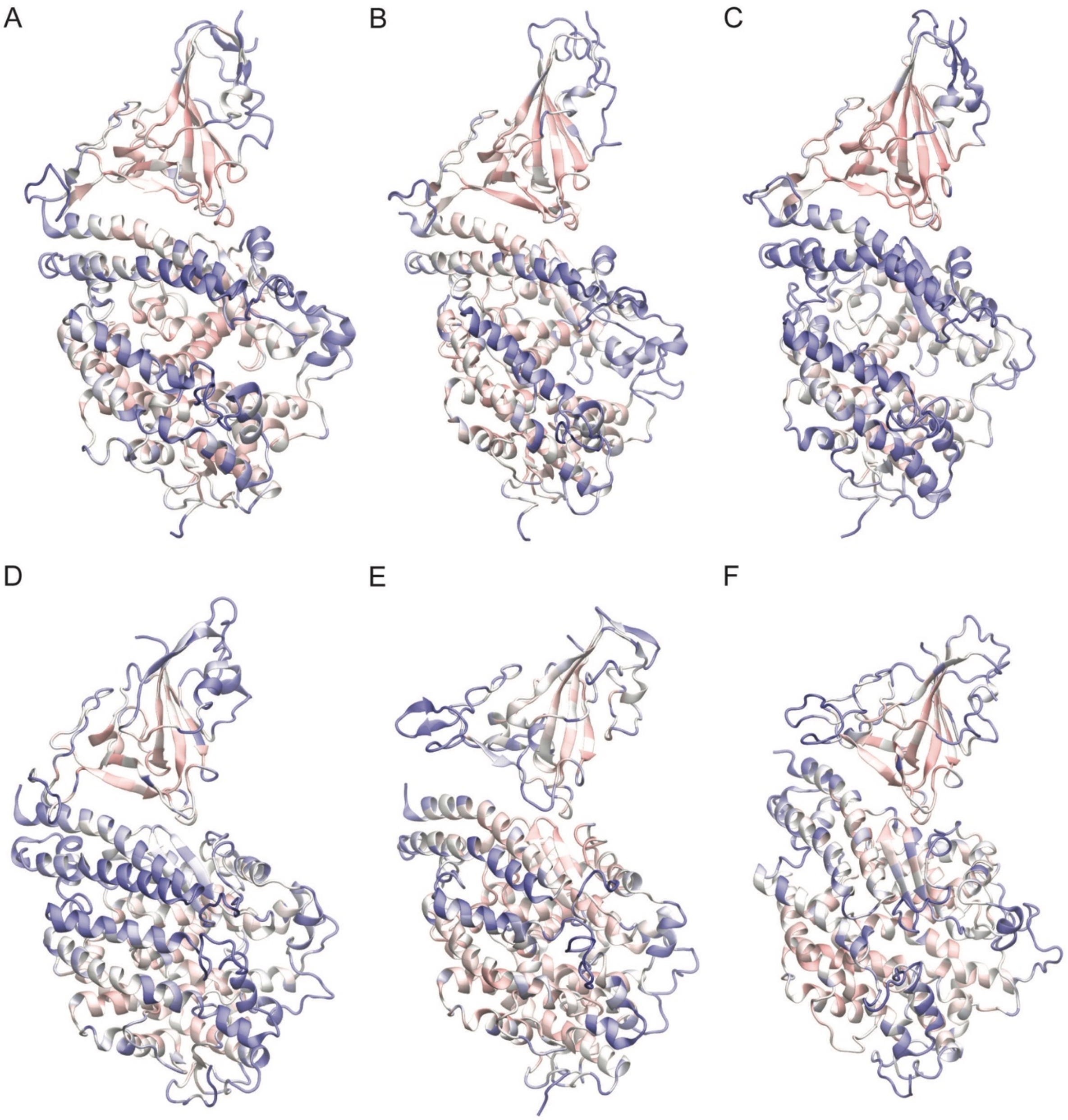


**Figure 2.** RMSF projected as beta factors to representative structures of SARS-CoV-2-ACE2 complex and SARS-CoV-ACE2. Pink shows regions of low flexibility and blue shows regions of high flexibility. A-C) SARS-CoV-2-ACE2 complex in three runs. D-F) SARS-CoV-ACE2 complex in three runs.

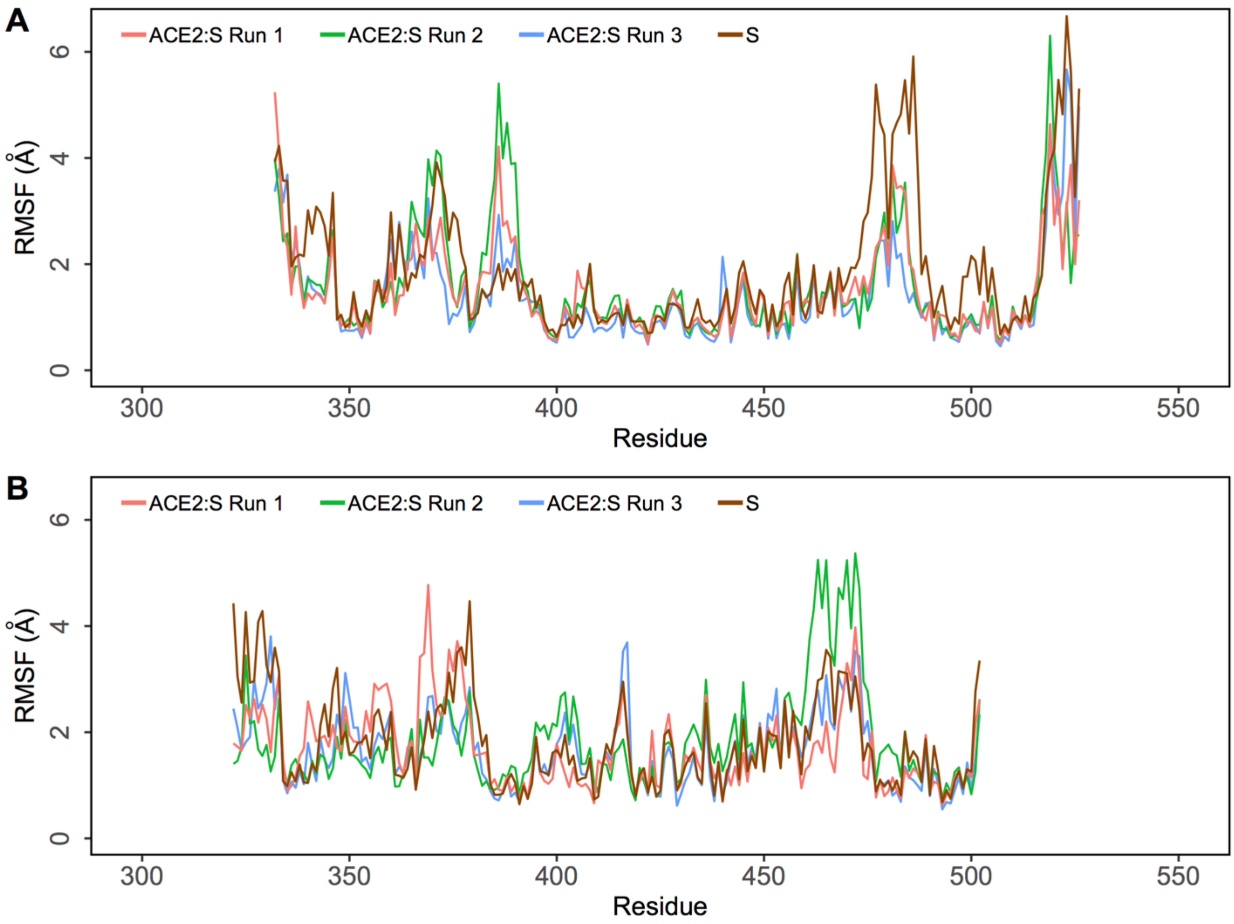


**Figure 3.** Root mean square fluctuation (RMSF) of protein Cα atoms obtained from three independent 500 ns runs of SARS-CoV-2 and SARS-CoV bound simulations of ACE2 compared to unbound simulations of SARS-CoV2 RBD and SARS-CoV RBD respectively. A) RMSF of Cα atoms of SARS-CoV-2 spike protein RBD in complex with ACE2 and without ACE2; B) RMSF of Cα atoms of SARS-CoV spike protein RBD in complex with ACE2 and without ACE2.

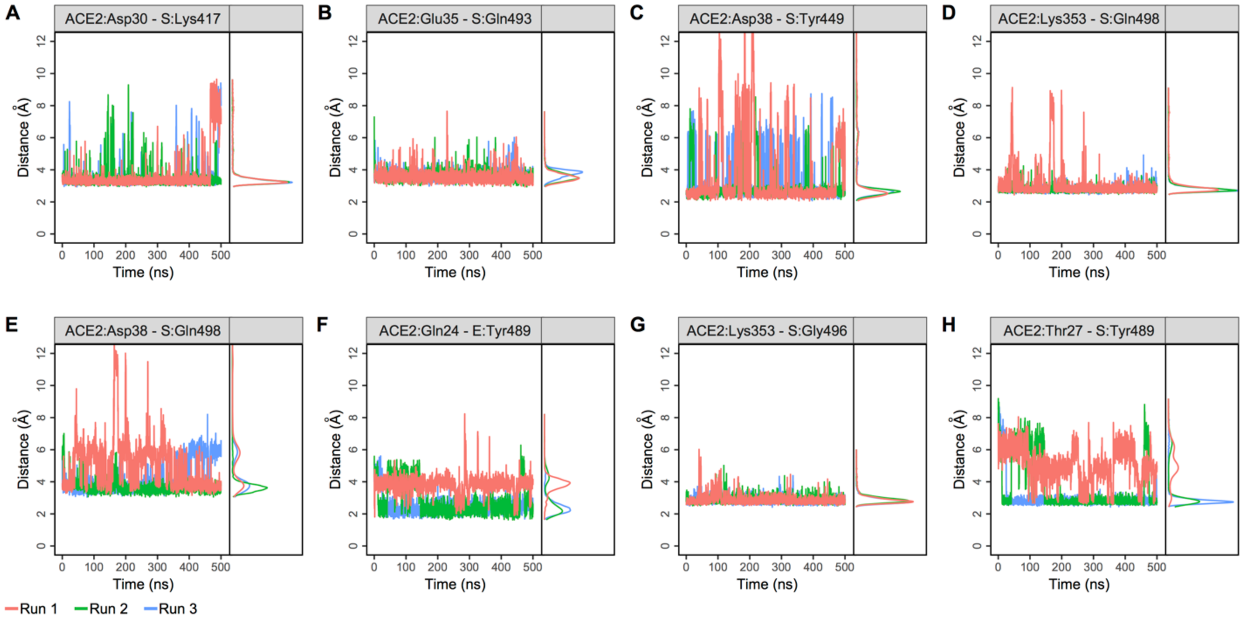


**Figure 4.** Contact distance plots of the most stable interfacial residues of SARS-CoV-2 spike RBD with ACE2. The contact distance was evaluated between specific atoms as indicated below to account for the possibility of switches between different hydrogen atoms (in Lys) or oxygen atoms (in Asp/Glu) that is involved in a hydrogen bond. Data from three runs are shown in red, green, and blue. The contact distance density plot of consistently interacting residues showed sharper peaks in the density plot adjacent to each distance plot. A) Distance between ACE2:Asp30_CG and S:Lys417_NZ; B) Distance between ACE2:Glu35_CD and S:Gln493_NE2; C) Distance between ACE2:Asp38_CG and S:Tyr449_HH; D) Distance between ACE2:Lys353_NZ and S:Gln498_OE1; E) Distance between ACE2:Asp38_CG and S:Gln498_NE2; F) Distance between ACE2:Gln24_O (backbone) and S:Tyr489_HH; G) Distance between ACE2:Lys353_NZ and S:Gly496_O (backbone); H) Distance between ACE2:Thr27_OG1 and S:Tyr489_OH.

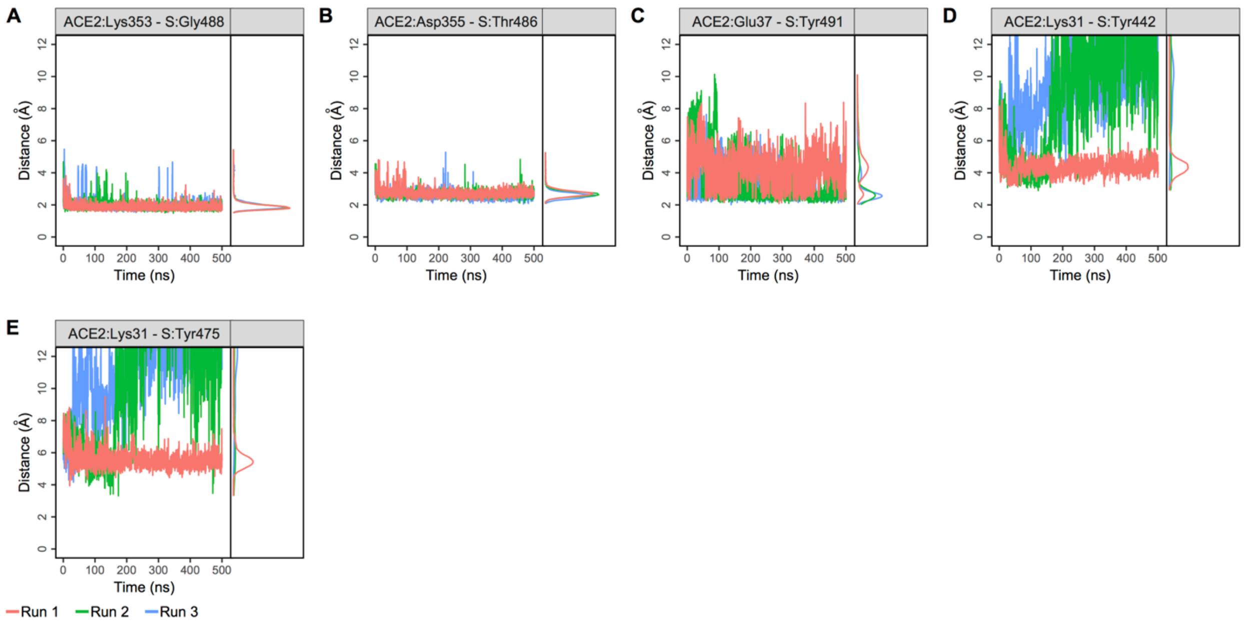


**Figure 5.** Contact distance plots of the most stable interfacial residues of SARS-CoV spike RBD with ACE2. The three runs are shown in red, green, and blue. The contact distance density plot of consistently interacting residues showed sharper peaks in the density plot adjacent to each distance plot. A) Distance between ACE2:Lys353_O (backbone) and S:Gly488_H (backbone); B) Distance between ACE2:Asp355_CG and S:Thr486_HG1; C) Distance between ACE2:Glu37_CD and S:Tyr491_HH; D) Distance between ACE2:Lys31_NZ and S:Tyr442_CZ (cation-π); E) Distance between ACE2:Lys31_NZ and S:Tyr475_CZ (cation-π).

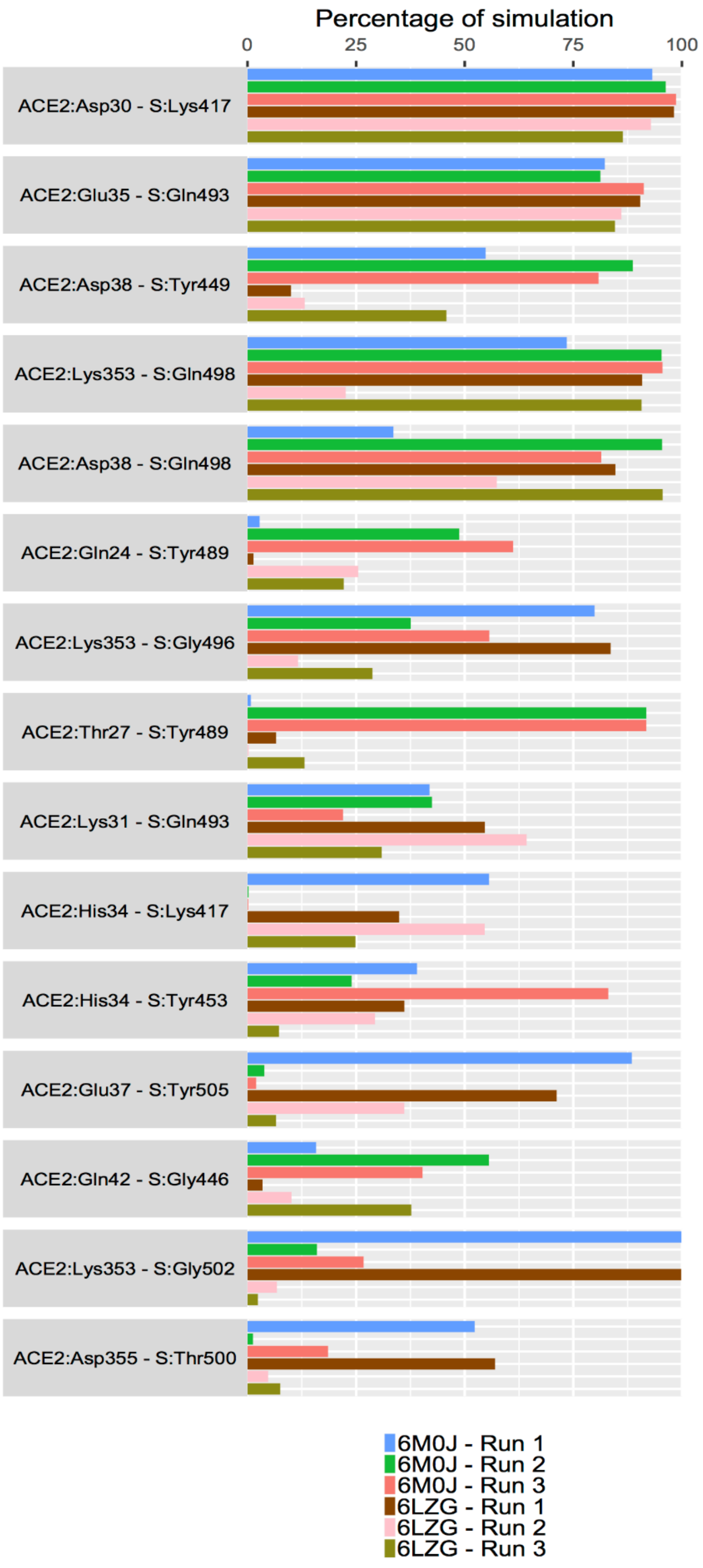


**Figure 6.** The percentage of simulation time during which specific intermolecular contacts were maintained between ACE2 and SARS-CoV-2 spike RBD residues in three independent runs each of structures with PDB IDs 6M0J and 6LZG.

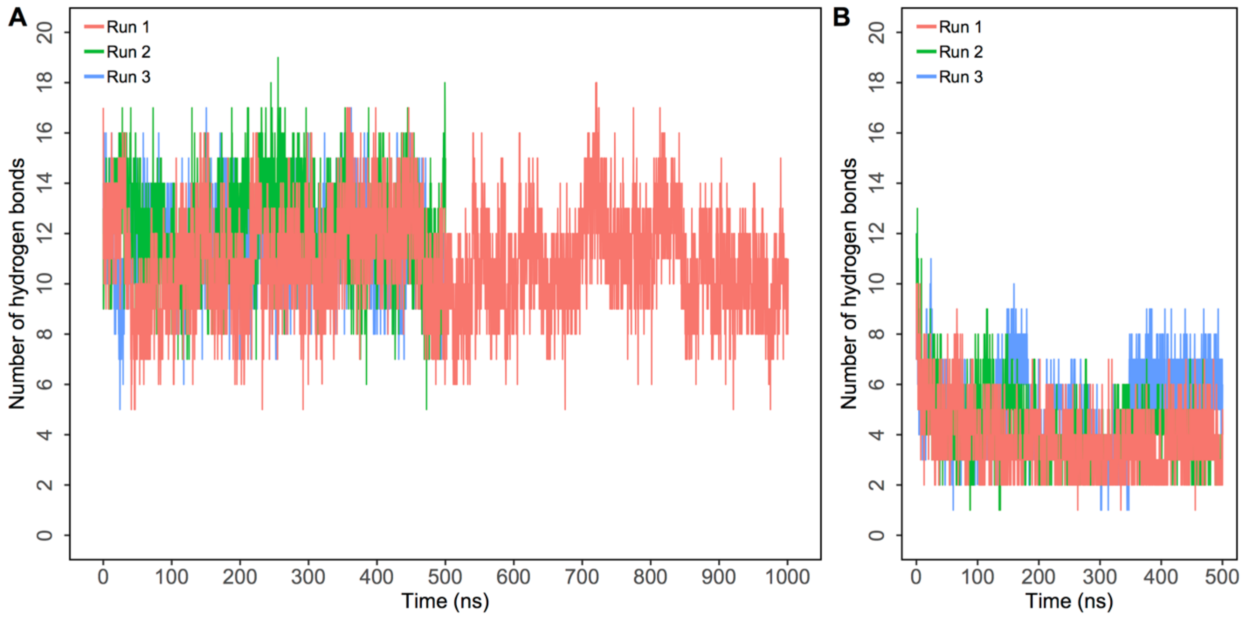


**Figure 7**. Number of intermolecular hydrogen bonds formed between SARS-CoV-2/SARS-CoV spike protein RBD and ACE2. Three independent runs are shown in red, green, and blue. A) Number of hydrogen bonds formed between SARS-CoV-2 spike RBD and ACE2; B) Number of hydrogen bonds formed between SARS-CoV and ACE2.
